# Coping switches stress–evoked network disconnection to a new forebrain network state

**DOI:** 10.1101/2024.08.19.607747

**Authors:** Laura Durieux, Karine Herbeaux, Nelson K Totah, Lucas Lecourtier

## Abstract

Coping with stress is critical to maintaining mental and physical health. Acute stress is associated with changes in neuronal activity across many brain regions and with the reorganization of cross– region correlations in neurovascular coupling. How neuronal activity itself is coordinated across regions during stress, and how neuronal networks are modified during coping, are unknown. We recorded in rats local field potentials (LFPs) simultaneously from five stress–responsive regions during stress and stress–coping behavior. We characterized network activity by computing cross– region coherence, Granger causality, and phase–amplitude coupling on bipolar derivatives of LFPs in the lateral habenula, basolateral amygdala, dorsal hippocampus, prelimbic cortex, and anterior cingulate cortex. First, we established a stress–coping model in rats. We showed that the behavioral response to acute 10–minute restraint stress returned to baseline after a second restraint 3 hours later. The pre–stress state was characterized by robust global network interactions in the theta (6– 9 Hz) and gamma (45–65 Hz) bands. Stress exposure led to nearly complete loss of connectivity. During coping, the robust connectivity reemerged, but in a new pattern compared to the pre–stress state. Finally, we found that baseline, stressed, and coping states can be predicted with high accuracy (> 90 %) from network activity. Overall, we showed that the acutely stressed brain state was primarily a state of network disconnection, while coping was a new network state rather than a return to baseline.

## Introduction

Stress is a major factor in mental and physical illnesses(Godoy et al., 2018). Coping with stress is an important aspect of daily life. Inadequate coping is associated with depression, anxiety, and post– traumatic stress disorder(Godoy et al., 2018),(Chrousos, 2009),(Nestler and Russo, 2024),(McEwen et al., 2015), so that it is of great importance to understand how the brain responds to stressful situations and how it adapts when coping with stress. Stress alters the activity of numerous brain regions involved in the perception of stress, the stress response, and adapting to stress, including the hypothalamic–pituitary–adrenal axis(de Kloet, 1992),(de Kloet et al., 1999), the noradrenergic system(Pace and Myers, 2023), the medial prefrontal cortex (mPFC)(McKlveen et al., 2015), the hippocampus (HPC)(Suri and Vaidya, 2015), and the amygdala (AMY)(Sharp, 2017).

Considerably less is known about the cross–region coordination of neuronal activity during stress and during coping. Prior studies have used Fos protein expression to resolve the network activity patterns in the context of stress(Knox et al., 2016),(Vetere et al., 2017),(Careaga et al., 2019),(Durieux et al., 2020) and shown that the lateral habenula (LHb), mPFC, AMY and HPC were functionally connected in response to stressor exposure(Durieux et al., 2020),(Durieux et al., 2022). However, Fos is an indirect measure of neuronal activity and is not expressed by all cell types(Salery et al., 2021),(Cruz-Mendoza et al., 2022). Most critically, Fos is a post–mortem measure relaying only a static snapshot rather than tracking the same network throughout stress and coping. Although fMRI–BOLD signals have yielded important insights into network activity in the context of stress and coping(Sinha et al., 2016),(Yuan et al., 2021),(Norbury et al., 2023), the neurovascular signal has a complex relationship with neuronal spiking, as well as slower changes in extracellular field potentials(Logothetis, 2003),(Ekstrom et al., 2009),(Magri et al., 2012). On the other hand, the local field potential (LFP), which is the superposition of the transmembrane currents produced (primarily) by synaptic inputs in the vicinity of the electrode, is ideal for inferring cross–region synaptic drive to define neuronal networks based on the underlying regional neuronal activity(Fries, 2005),(Buzsáki et al., 2012). To our knowledge, LFP recordings have not been made simultaneously across numerous stress–responsive regions, neither during the response to stress, nor during coping.

Here, we investigated the cross–region coordination of neuronal activity among the LHb, dorsal HPC (dHPC), basolateral AMY (BLA), prelimbic cortex (PRL), and anterior cingulate cortex (ACC) in male rats. In the context of repeated stress exposures (two 10–minute restraint sessions 3 hours apart), we recorded multi–electrode LFPs within each of these five regions, simultaneously, and used local bipolar derivations in each region for computing cross–region coherence, Granger causality, and phase–amplitude coupling. We used a variety of behavioral measures to demonstrate coping after the second stressor. We characterized how cross–region neuronal interactions change in response to stress and during coping, focusing on the active wake period, when attention and information processing are at the highest following an emotionally salient event(Fries, 2023),(Karakaş, 2020),(Okonogi and Sasaki, 2021).

## Results

We aimed to characterize neuronal activity coordination across multiple stress–responsive regions, during the acute response to stress and during coping to a repeated stressor. We performed a repeated stress procedure (n = 10 rats) that consisted of a baseline (BSL) session on the preceding day followed by two 10–minute restraint 3 hours apart (**Fig. 1A**). Behavior was scored and LFPs were recorded. LFPs are the superposition of the transmembrane currents produced (primarily) by synaptic inputs in the vicinity of the electrode, which makes the LFPs ideal for inferring synaptic drive between brain regions(Logothetis, 2003),(Fries, 2005),(Buzsáki et al., 2012). We calculated bipolar derivations from recording sites in five stress–responsive brain regions, simultaneously (**Fig. 1B-C**). Analyses were done on data collected during the active wakefulness (see supplementary methods for vigilance classification).

**Fig.1.**
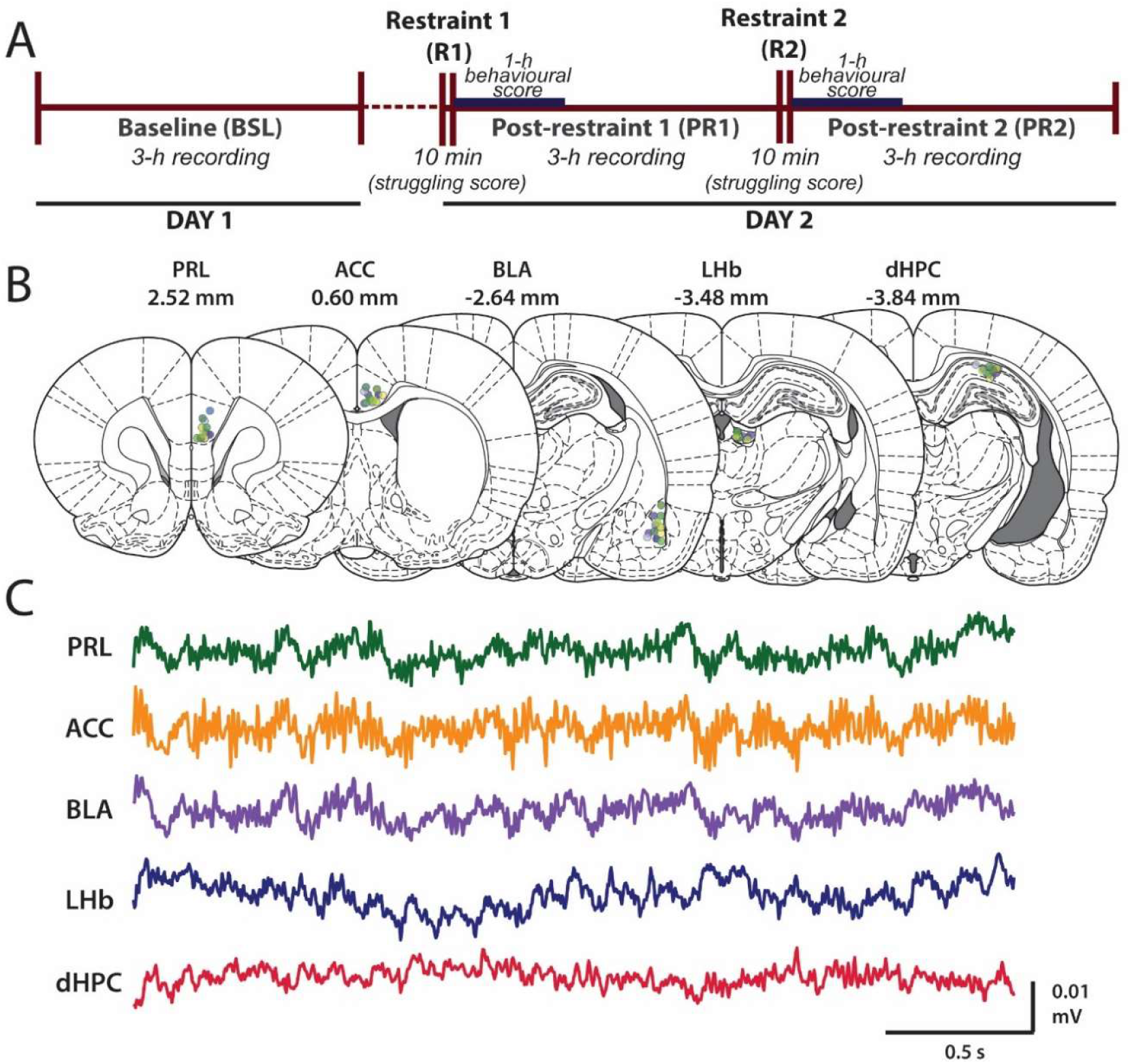
We recorded LFPs from 5 forebrain regions simultaneously during a repeated restraint stress paradigm. **A**) The timeline of the experiment is shown. On day 1, we made a 1–hour baseline LFP recording and quantification of behavior. The stress exposure experiment took place on the following day (day 2). It consisted of two 10–minute restraint stressors, during which the time spent struggling was recorded. Each stressor was followed by a 3–hour recording during which LFPs were recorded and behaviors scored during the first hour; these are referred to as the post–restraint 1 (PR1) and post–restraint 2 (PR2) epochs of the experiment. **B**) Post–mortem histology was used to determine the locations of the 3–electrode bundles implanted within the PRL, ACC, BLA, LHb, and the CA1 region of the dHPC. The site for each rat (n = 10) is shown in a different color. The numbers above each coronal brain diagram show the distance from Bregma in the anteroposterior axis. **C**) Representative filtered (0.5–100 Hz) voltage traces simultaneously recorded in the five regions of interest during the baseline.

### Rats exhibit multiple behavioral signs of coping to stress

First, we needed to establish a behavioral paradigm in which rats would show stress coping. We defined coping as a “normalization” or “return–to–baseline” of a variety of stress–evoked behaviors, including struggling during restraint, and post–restraint stress–related behaviors (locomotion, sniff down, digging in the cage bedding, wet–dog shakes, and circling). Rats spent less time struggling during the second restraint (7.18 ± 2.82 sec) compared to the first restraint (24.01 ± 7.52 sec) (Wilcoxon Signed–Rank Test, W = 2.80, p = 0.0051; **Fig. 2A**). The overall behavioral activation significantly differed across conditions (ANOVA, F(2,18) = 28.22, p = 0.0001). Post hoc tests indicated behavioral activation during the first post–restraint epoch (PR1) relative to baseline (p = 0.0003), but a return to BSL level following the second restraint (PR2) (p = 0.973 vs BSL and p = 0.0001 vs PR1) (**Fig. 2B**). Analysis of individual behaviors indicated that the first restraint preferentially impacts locomotion, sniff down, and digging: *Locomotion:* F(2,18) = 12.70, p = 0.0016; post hoc: BSL vs PR1, p = 0.033; PR1 vs PR2, p = 0.001; BSL vs PR2, p = 0.547. *Sniff down:* F(2,18) = 7.63, p = 0.0088; post hoc: BSL vs PR1, p = 0.038; PR1 vs PR2, p = 0.017; BSL vs PR2, p = 0.671. *Digging:* F(2,18) = 15.85, p = 0.0002; post hoc: BSL vs PR1, p = 0.0009; PR1 vs PR2, p = 0.023; BSL vs PR2, p = 0.143. *Wet–dog shake:* F(2,18) = 4.77, p = 0.00235; post hoc: BSL vs PR1, p = 0.152; PR1 vs PR2, p = 0.0504; BSL vs PR2, p = 0.496. *Circling:* F(2,18) = 2.15, p = 0.159; post hoc: BSL vs PR1, p = 0.944; PR1 vs PR2, p = 0.078; BSL vs PR2, p = 0.2649 (**Fig. 2C**). The paradigm used here likely results in the succession of two distinct stress states. During PR1, the large increase in the number of occurrences of stress– related behaviors indicates that this epoch represents a “stressed state.” This is supported by previous studies in rats showing the presence of physiological hallmarks of stress following restraint(Durieux et al., 2022),(Ngampramuan et al., 2008),(Wongsaengchan et al., 2023),(Jackson and Moghaddam, 2004),(Swanson et al., 2004),(Mo et al., 2008). On the other hand, rats struggle less during the second restraint and commit less stress–related behaviors in its aftermath. Thus, this epoch represents a “coping state.”

**Fig. 2.**
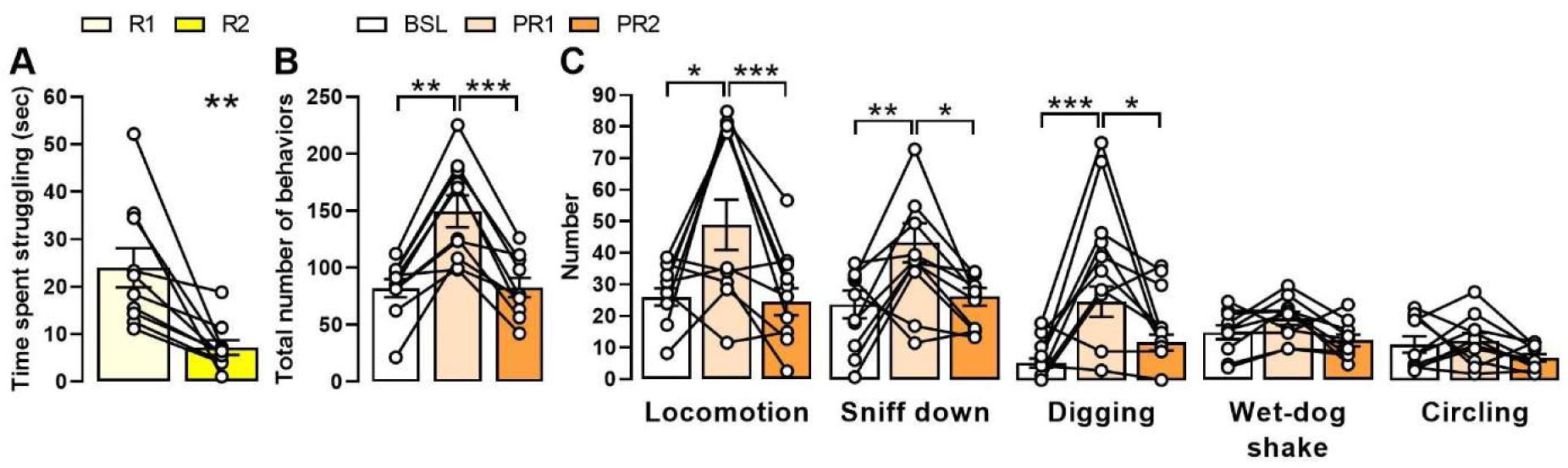
Struggling during restraint and stress–related behaviors after restraint both return–to–baseline after a second stressor. **A**) Struggling time during the first (R1) and second (R2) restraint (n = 10; Mean +/- SEM). We counted the number of times a given stress–related behavior was initiated during the first hour of the 3– hour post–restraint window 1 (PR1) and 2 (PR2) and compared them to the 1–hour baseline scoring on the prior day. **B**) Total number of stress–related behaviors (summed across all of the quantified behaviors) during the BSL, PR1, or PR2 (n = 10; Mean +/- SEM). **C**) The number of times each of the quantified behaviors was initiated during BSL, PR1, or PR2 (n = 10; Mean +/- SEM). White circles represent individual rats. *p < 0.05, **p < 0.01, ***p < 0.001.

### Stress decouples large–scale, global network activity patterns in the theta and gamma bands

Coherence between recording sites can be indicative of direct synaptic interactions, or common synaptic input to the two regions. We analyzed the bipolar derivatives at each recording site during the first minute of active wakefulness within each experimental condition because we wanted to capture the initial, most apparent and clear state proximal to the stress event. Spectral power analysis revealed theta and gamma band oscillations at all of the recording sites (**Supplementary Fig. 1**). We calculated cross–region coherence and tested for significance using bootstrapping and a two–sided permutation test. We found that coherence across all recording site–pairs is primarily in the theta and/or gamma band during both the BSL (**Fig. 3A**) and the PR1 epoch (**Fig. 3B**). Notably, the pairwise coherence is diverse across recording site–pairs showing that the LFP is not global (and not volume conducted) but is rather distinct for each cross–regional pair. We used graph theory to plot the topology of the stress network by drawing a link between recording sites if at least 80 % of the rats had significant theta (**Fig. 3C**) or gamma (**Fig. 3D**) coherence. Our results show that, relative to the BSL condition, acute stress exposure is associated with amygdalo–habenular and frontal– hippocampal disconnections in a network characterized by theta band oscillations. On the other hand, in the gamma band, the dHPC is completely segregated from the other four regions. The LHb also becomes disconnected from the ACC. Overall, our results suggest that there is widespread coherence between all five regions during the BSL state and that the stressed state is primarily characterized by widespread disconnection of many nodes of this BSL state network.

**Fig. 3.**
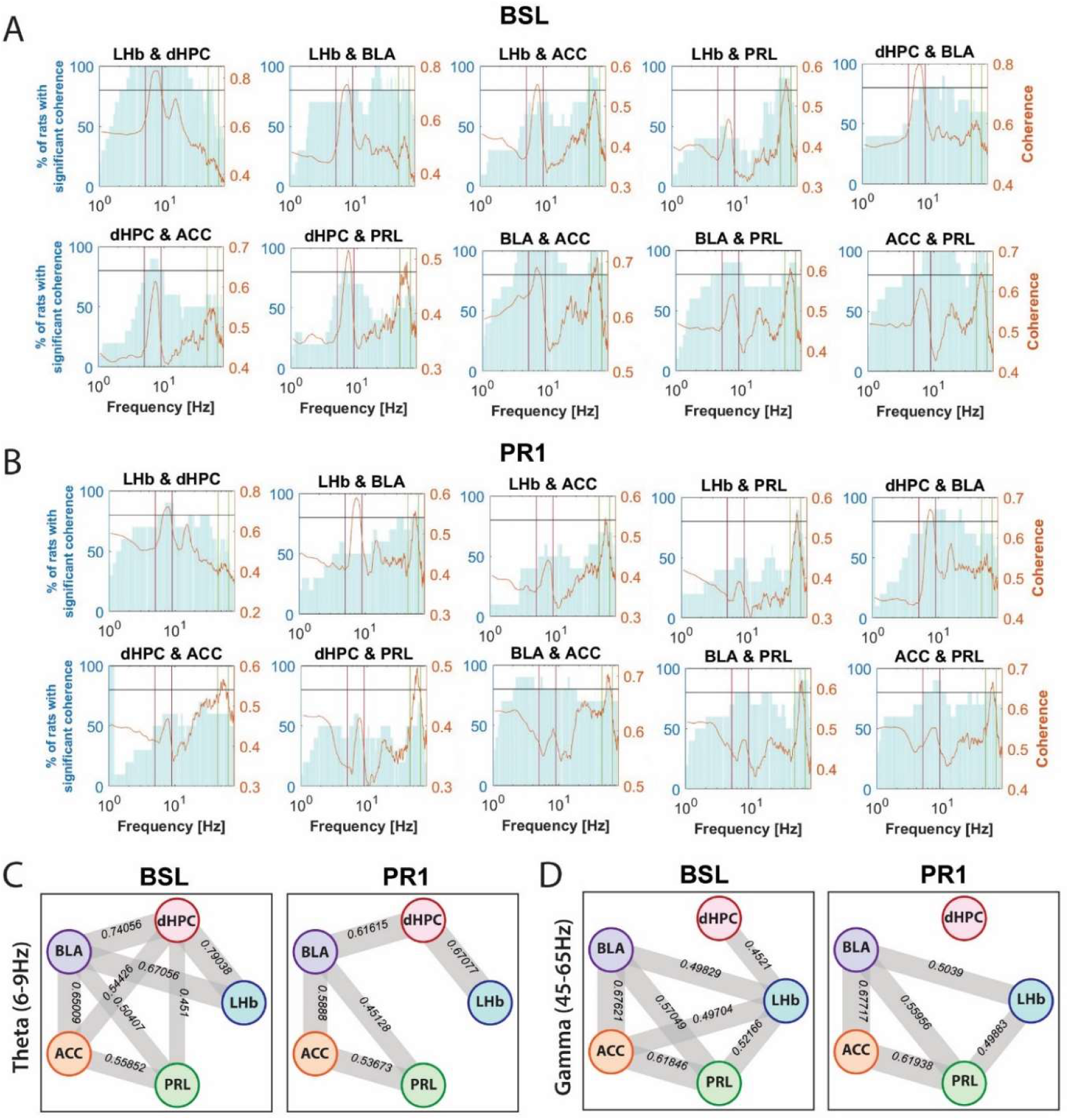
Acute stress response is associated with both theta and gamma band network disconnection. (**A–B**) Percentage of rats (left y–axis, blue bar plots) with significant coherency for each recording site–pair during BSL (**A**) and PR1 (**B**) conditions (n = 10 rats). The mean coherence (right y–axis, orange lines) is plotted across frequency (x–axis; logarithmic scale) between each of the 10 recording site–pairs. On each plot, the two vertical red lines frame the theta band (6–9 Hz), whereas the two vertical green lines frame the gamma band (45–65 Hz). (**C–D**) Graph networks depict significant cross–region coherence as thick grey lines. Line thickness is related to coherency magnitude. Panels C and D show the networks based on theta and gamma coherence, respectively.

### Coping with stress is associated with a novel network state

We have shown that the stressed state is associated with a loss of the widespread cross–region coherence observed during the pre–stress BSL. It is possible that coping behavior is associated with a restoration of this connectivity. We tested this hypothesis by assessing coherence during PR2. Again, we observed theta and gamma band coherence across many recording site–pairs (**Fig. 4A**). It is apparent that, in both the theta (**Fig. 4B**, left) and gamma (**Fig. 4B**, right) bands, the network activity does not return to the BSL state. There are a few notable characteristics of the network during the coping state. In the theta band, there is little change from PR1, other than the additional loss of ACC–PRL connectivity. In the gamma band, the network is also highly similar to PR1, except with the addition of a dHPC–BLA functional connection. Thus, stress coping behavior is associated with modification of the stressed (PR1) state, with both gains and losses of cross–region coherence, as opposed to a restoration of the original BSL state–network pattern.

**Fig. 4.**
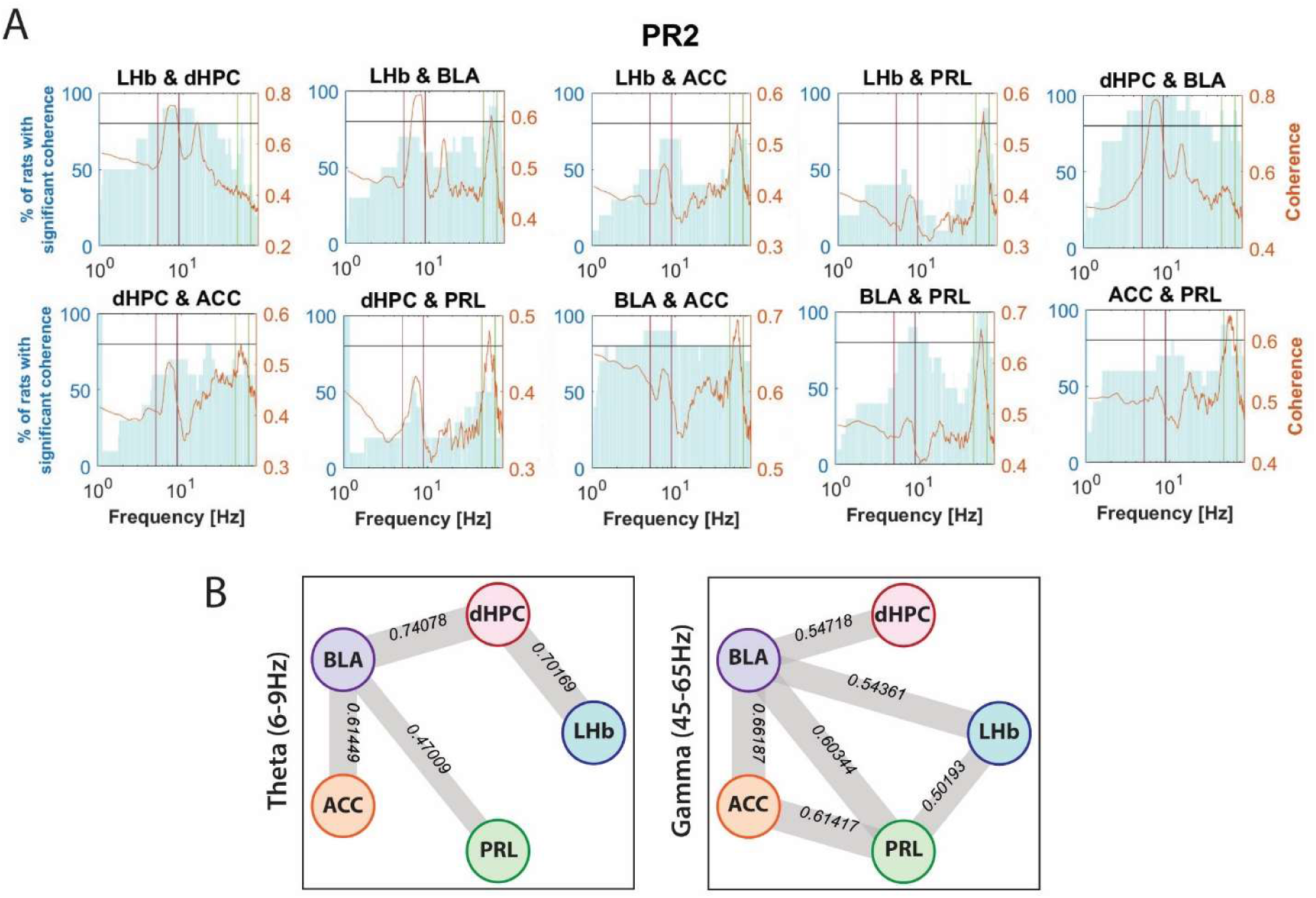
Coping is accompanied by the emergence of a novel network connectivity pattern compared. **A**) Percentage of rats (left y–axis, blue bar plots) with significant coherency for each recording site–pair during the PR2 condition (n = 10 rats). The mean coherence (right y–axis, orange lines) is plotted across frequency (x–axis; logarithmic scale) between each of the 10 recording site–pairs. On each plot, the two vertical red lines frame the theta band (6–9 Hz), whereas the two vertical green lines frame the gamma band (45–65 Hz). **B**) Graph networks depict significant cross–region coherence as thick grey lines. Line thickness is related to coherency magnitude. The left and right panels show the networks based on theta and gamma coherence, respectively.

### Shared synaptic input does not explain network changes during stress or coping

The LFP signals in a pair of brain regions may be coherent due to synaptic inputs that are common to the recorded regions. Although global fluctuations in brain activity are mitigated by using the bipolar derivations of the LFP signals, as we have done above, we also calculated site–pair Granger causality magnitudes as a measure of directional causality that accounts for common synaptic input by conditioning pairwise Granger causality on all other regions in the network. **Figure 5** shows that not all recording sites with significant coherence are associated with directional interactions. However, in agreement with the coherence analysis, the BSL condition is associated with large–scale uni– and bi–directional interactions in the theta and gamma bands, while the stressed state (PR1) is associated with large–scale disconnection. In the theta band, out of the five recording site–pairs with directional interactions observed in the BSL epoch, only two are spared from disconnection (dHPC–BLA and dHPC–LHb). In the gamma band, this effect is even more robust with a total loss of network connectivity.

**Fig. 5.**
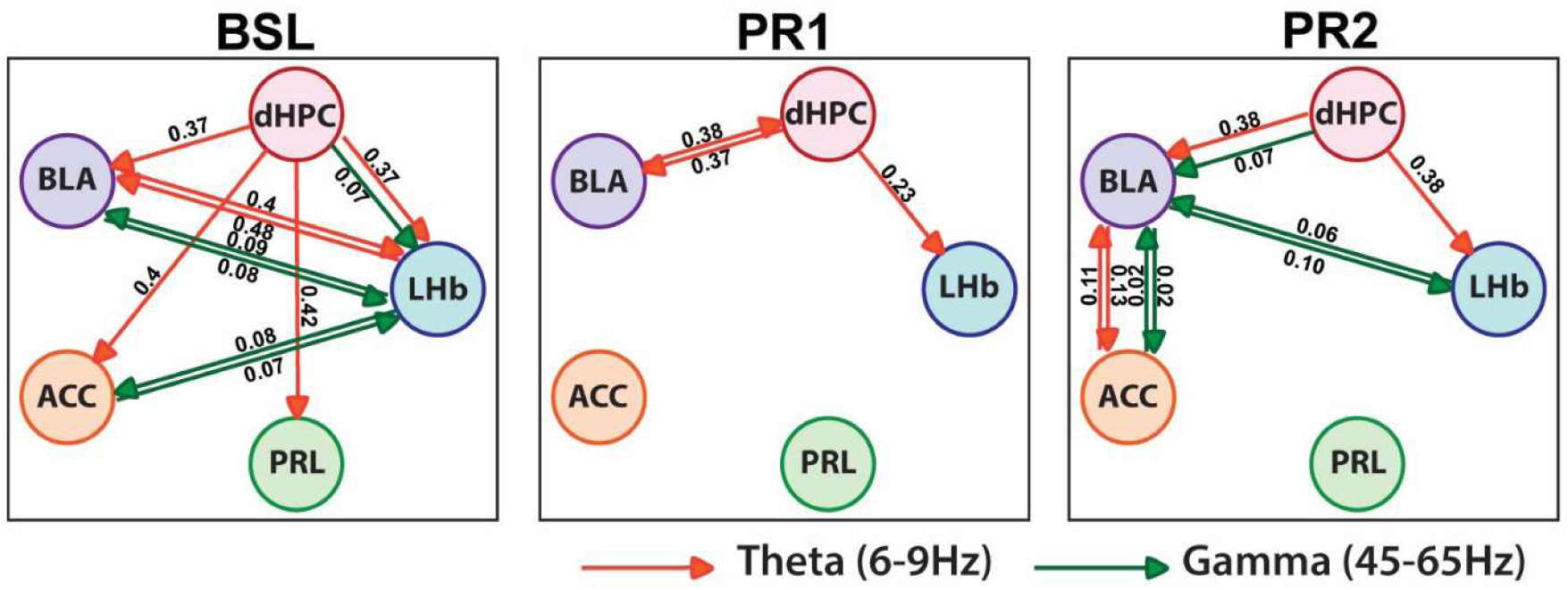
Granger causality analysis revealed a stress–associated state of network disconnection and a coping state characterized by a new network connectivity pattern. Graph networks depict relationships within the network in the theta (*upper row*) and gamma (*lower row*) bands for BSL (*left*), PR1 (*middle*), and PR2 (*right*) sessions. The values above the lines indicate granger causality magnitude (only significant values shown).

During coping (PR2), the network does not return to the pre–stressed state, nor does it remain in the PR1 state. Instead, a gain of connectivity occurs and produces a distinct and new network state relative to the PR1 and the BSL states. In the theta band, this new network is characterized by the emergence of an ACC–BLA bidirectional connection that occurs in neither the pre–stress or stressed states. In the gamma band, this same connection also appears during coping. Moreover, the total connectivity loss during PR1 is reverted with the restoration of one pre–stress pattern (LHb–BLA bidirectional interaction) and the emergence of two new patterns in the gamma band (dHPC–to– BLA and bidirectional ACC–BLA connections).

Overall, these analyses suggest that a first stressful experience is associated with a general disconnection between the mPFC and related regions, and between the BLA and LHb, with the preservation of bidirectional connection between the dHPC and BLA, and dHPC to LHb connection. Coping is associated with the restoration of the bidirectional connection between the BLA and both the ACC (theta and gamma jamais partie vs cohérence) and LHb (theta only) as well as a novel unidirectional theta band connection between dHPC and BLA.

### Cross–frequency interactions between different regions are highly diverse

The presence of both theta and gamma band directional interactions suggests the possibility that these temporal patterns of synaptic activity may interact. Determining whether such cross– frequency interactions occur is important for generating mechanistic predictions about the cross– region drive of specific cell types or transmembrane currents that generate slower (theta band) and faster (gamma band) field potential fluctuations. We calculated the degree to which gamma band oscillation amplitude in one region had a stable pattern of temporal modulation by theta band oscillation phase in the other region(Tort et al., 2010a). **Figure 6** shows the modulation index magnitude for recording site–pairs that had significant site–pair Granger causality. Strikingly, we observed some regions largely devoid of cross–region theta phase modulation of gamma power (plots which are mostly black). Moreover, we observed instances of differential modulation of low gamma power (< 60 Hz) and high gamma power (> 60 Hz). Finally, for some cross–region pairs, modulations dependended differently on theta phase in each region. This analysis demonstrates that there are highly diverse cross–region, cross–frequency interactions in the stress and coping networks.

**Fig. 6.**
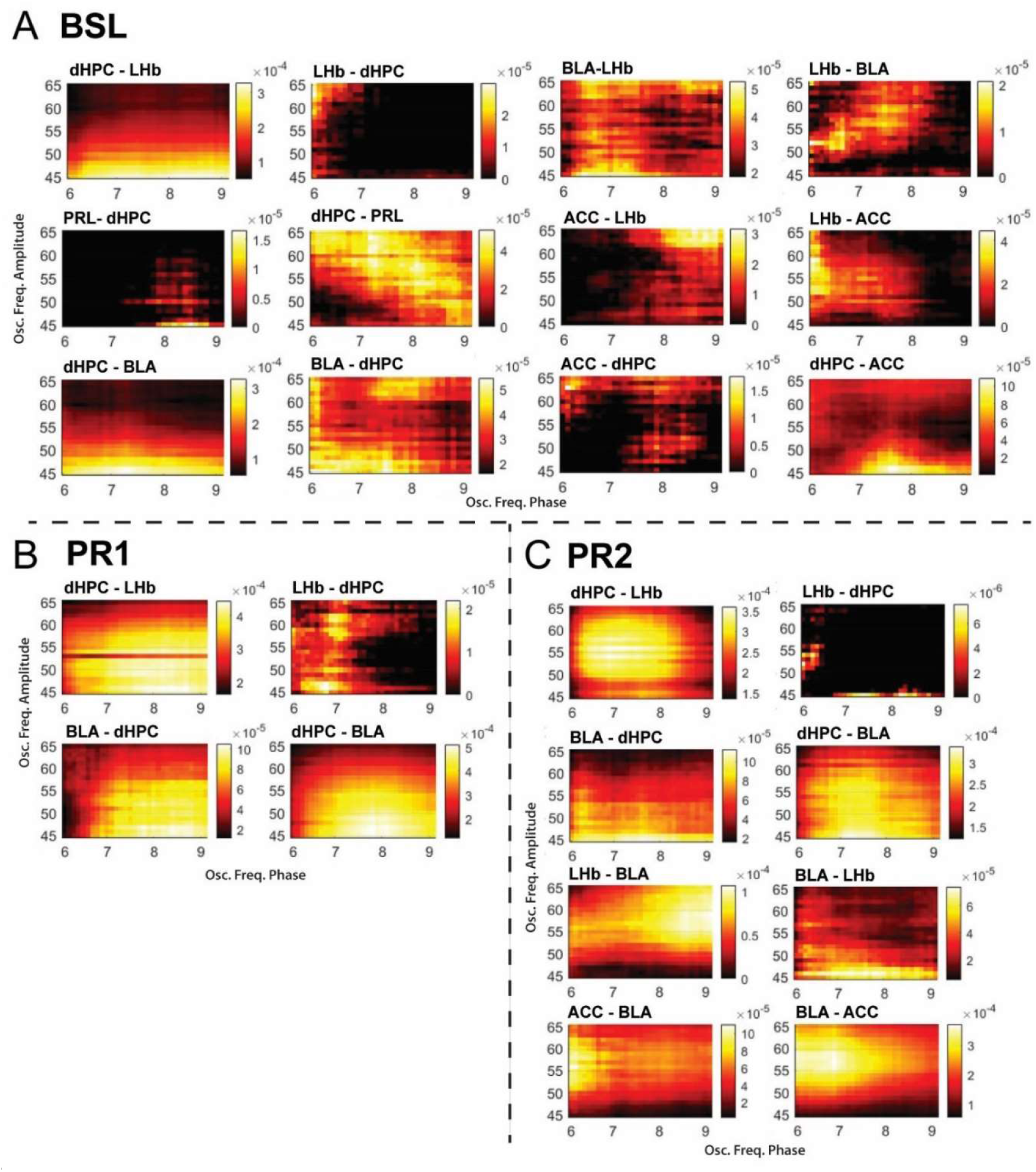
The stress and coping networks are associated with diverse cross–region, cross–frequency phase– amplitude coupling patterns. We calculated a modulation index, which was statistically corrected so that only those values above zero (not black) are significant. The first region listed in title of each plot was used for theta phase and the second region was used for gamma amplitude. The x–axis spans a broad theta band and the y–axis spans a wide gamma band, which enables visualization of changes in subregions of these two frequency bands. The panels show the modulation index between recording site–pairs with significant coherence during the baseline condition (**A**), the PR1 condition (**B**), and the PR2 condition (**C**).

### Stress and coping states can be decoded from network activity

We observed distinct neuronal network activity associated with pre–stress, stressed, and coping states. These neuronal network states are a potential biomarker for stress and coping. Given that we have recorded field potentials, such a biomarker would have translational potential for human field potential recordings using intracranial EEG. We sought to decode the pre–stress, stressed, and coping states from the coherence magnitudes recorded across the network. We chose coherence, over Granger causality, because coherence can be calculated online as a low–latency biomarker. We trained a feed–forward neural network and tested it on held–out data. The experiment condition (baseline, PR1, or PR2) could be predicted from network coherence patterns with an average accuracy of 91.7% (**Fig. 7**, p<0.001). Prediction was also successfully achieved when 9 out of the 10 rats were used for training and the prediction made with data of the remaining rat (**Supplementary Fig. 3**). Therefore, coherence across this network holds potential as a biomarker for tracking stress and coping.

**Fig. 7.**
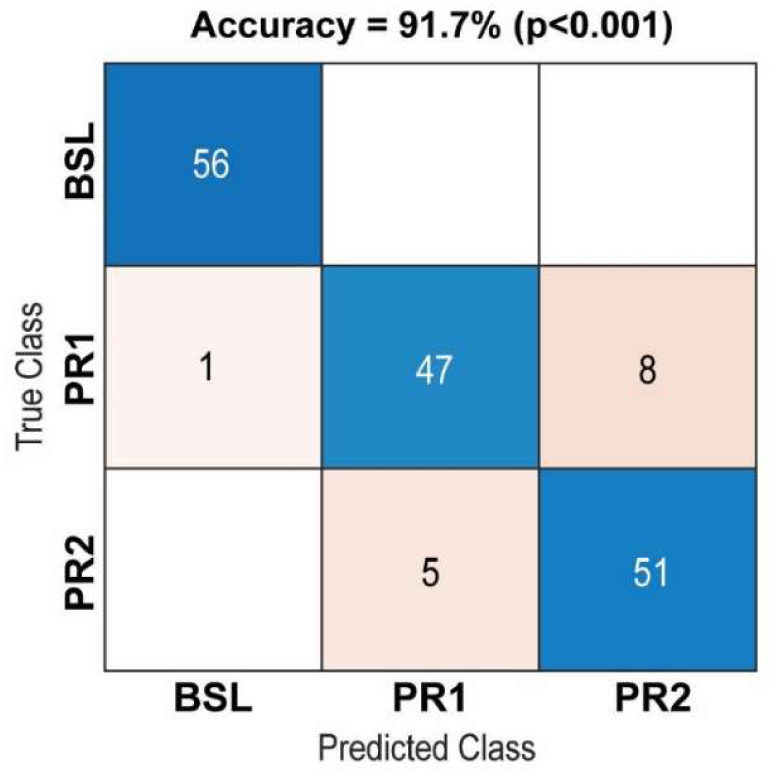
Experiment conditions can be predicted from network coherence patterns. The confusion chart shows prediction accuracyThe blue squares represent cases when the algorithm returned the correct prediction and the red squares represent the wrong predictions. The number of observations are written in each square.

## Discussion

Here, we report that distinct neuronal network states are associated with baseline, stressed, and coping states. The baseline state was characterized by robust global connectivity, the stressed state by sparse connectivity, and the coping state by the reemergence of robust connectivity in a novel pattern compared to baseline. We show that behavioral state can be decoded from the network state. In interpreting our results, it is important to note that the LFP is dependent upon the orientation of the surrounding neurons and the cytoarchitecture of some subcortical regions poses difficulty in interpreting the LFP. Nevertheless, these recordings show the dynamics of synaptic inputs coordinated across a network of forebrain regions, which has heretofore relied on indirect measures neuronal metabolism and neuronal activation garnered from slow, hemodynamic signals.

### Pre–stress baseline is associated with widespread network connectivity

While it is known that the PFC, HPC, and AMY have coordinated activity in awake rats(Jacinto et al., 2013),(Stubbendorff et al., 2023), our findings suggest that the LHb is also engaged in this forebrain network. For instance, we observed LHb–dHPC theta coherence. This pattern has been observed in awake rats and in the context of cognitive functions(Aizawa et al., 2013),(Goutagny et al., 2013). We also observed coherence between the LHb and the BLA, as well as the mPFC, in in line with prior studies using Fos, a static indirect measure(Durieux et al., 2020),(Durieux et al., 2022). Overall, the present findings indicate widespread forebrain connectivity, including areas such as the LHb.

### Stress produces widespread network disconnection

Animal studies of the neurophysiological consequences of stress exposure have been primarily in the domain of early life stress or chronic stress, and there is paucity of functional connectivity data concerning acute stress and subsequently coping with repetitions of the stressor. Decreased BLA–mPFC theta coherence has been reported in rats after maternal separation(Cao et al., 2016) and after chronic exposure to an elevated platform(Lee et al., 2011). We observed no change. Gamma coherence has been shown to increase between dHPC and PRL/ACC as well as between ACC and PRL after chronic social defeat stress. We observed no change in this aspect of the network either. Therefore, we conclude that the brain state evoked by the acute stress paradigm used in the present study may differ from the brain state formed during chronic stress.

Our study involves regions related to the attention/cognitive control and threat/salience circuitries, as proposed by Nestler and Russo(Nestler and Russo, 2024), and implicates specifically the theta and gamma bands in the neuronal network state during acute stress. This finding is in line with prior work on how neuronal activity is coordinated between two brain regions during acute stress(Colgin, 2013),(Headley and Paré, 2013). For instance, during fear conditioning, the AMY and HPC, as well as the AMY and PFC, have coherent LFP signals in these bands(Seidenbecher et al., 2003),(Likhtik et al., 2014). Moreover, similar findings have been reported between the PFC and HPC in an anxiogenic context(Adhikari et al., 2010).

### Neuronal network activity may provide a biomarker for assessing stressed and coping states

We observed that the stressed state was a sparse network and the coping state was characterized by robust connectivity. These network states were compared to a pre–stress baseline (which itself had robust connectivity, but in a different pattern than the coping state). However, it is possible that the state following the second stressor was a transition state and that, with adequate time, the network might relax into to baseline state. Similarly, the baseline state network configuration may itself be dynamic. Future work can address these questions using multi–day pre–stress baseline recordings and continuous (24 hours per day) recordings after the second stressor. If the pre–stress, stressed, and coping states are associated with stable forebrain networks, then our findings hold potential as a biomarker for stress and coping. Consistent with this potential, we were able to successfully predict with high accuracy (> 90 %) the experimental condition from the forebrain network activity. Biomarkers for stress and coping have typically used peripheral measures that correlate with activity of the autonomic nervous system(Carnevali and Sgoifo, 2014), or circulating corticosterone(Armario et al., 2023). Our results suggest that monitoring field potentials may hold promise for tracking stress and coping. Neuromodulation therapies aimed at steering neuronal networks toward a state of coping may also need to consider that coping is a new brain state, rather than a return to the “normal”, pre–stress baseline state.

In sum, we observed specific patterns of neuronal network activity between regions in a cortical– hippocampal–amygdalar–habenular network. There was a stark contrast in network activity between the pre–stress and post–stress states. Yet, a new network state emerged during coping that resembled neither the baseline nor the post–stress state. These finding suggests that treating stress–related disease may require modulating neuronal networks into a new state, rather “normalizing” the brain to its original state prior to stress. The ability to predict pre–stress, stress, and coping states from the neuronal network state holds potential usefulness as a biomarker in tracking and treating stress–related disease.

## Methods

### Animals

Experimental protocols and animal care were in compliance with the institutional guidelines (council directive 87/848, October 19, 1987, Ministère de l′agriculture et de la Forêt, Service Veterinaires de la Santé et de la Protection Animale) and international laws (directive 2010/63/UE, February 13, 2013, European Community) and policies: APAFIS#7114. The study required 15 male Long–Evans rats (250–350 g; Janvier Labs, France). Rats were pair housed (until surgery) on a 12 hour light cycle (light at 07:00) at 22 ± 1 °C (∼55 % RH) with unrestricted food and water access.

### Surgery

Under gas anesthesia, a bundle of three LFP recording electrodes (tungsten) was implanted in each region of interest: PRL, ACC, BLA, LHb, and dHPC (see **Supplementary Information** for stereotaxic coordinates and detailed implantation procedure).

### Stress paradigm

Two 10–minute restraint sessions were performed 3 hours apart (**Fig. 1A**). See **Supplementary Information** for detailed procedure.

### Assessment of behavior

During both restraints, the time spent struggling was manually scored by the person doing the restraint. Multiple stress–related behaviorswere quantified from video analysis using an in–house detection algorithm (courtesy of Elouan Cosquer) during the first hour of the baseline and during the first hour following each restraint session. These behaviors were: *locomotion* (rats moving from an immobile state, or walking from one area of the cage to another), *sniff down* (rats positioned head down with their nose below the horizontal plane of their body and sniffing, while either standing or walking)(Cole and Koob, 1989), *digging* (spreading the sawdust with the forepaws), *wet*– *dog shake* (paroxystic shudder of the head, neck, and trunk)(Bedard and Pycock, 1977), and *circling* (making a complete turn in a tight space)(Costall and Naylor, 1974). The number of occurrences of each behavior was counted.

### Electrophysiology signal processing and analyses

#### Signal acquisition

The broadband (0.5 Hz to 8 kHz) signal was sampled at 1,375 Hz (AlphaLab amplifier, Alpha Omega, Germany; and a preamplifier from AD Instruments, UK). Electrode impedance at 1 kHz was ∼ 50 to 200 kΩ.

#### Pre–processing

Analyses were done in MATLAB R2020b (MathWorks). Signals were downsampled to 275 Hz and then bandpass filtered (0.5–100 Hz) using a 2^nd^ order Butterworth acausal filter. To avoid contamination of the local signal by volume–conducted signals from surrounding regions, for each region, we calculated bipolar derivations by subtracting the signals recorded on two of the three channels. We chose the two electrodes with minimal artifactual contamination throughout the experiment. We then removed slow drift using the locdetrend function from the Chronux toolbox (http://chronux.org/). Movement artifacts were removed using an artifact detection and removal algorithm based on stationary wavelet transform(Islam et al., 2014). Signals were visually inspected to ensure that artifacts and clipped signals were excluded from analysis.

#### Spectral analyses

All analyses used the Chronux toolbox. Power was calculated using a 4–second sliding window in 0.2 second steps. Tapers were set to k = 9 and nw = 5. Power was Z–scored, within each frequency bin, to the time–binned power of the recorded signal. Coherence was calculated for each cross–region recording site–pair, on the first minute of the awake state occurrence in each condition (baseline, stress, or coping). We assessed whether coherence values were significant using a one–sided permutation test (p < 0.01). A distribution of 1,000 chance coherence values was generated by bootstrapping. Surrogate data were made by cutting the signal at a random time point and flipping the two signal segments in time(Canolty et al., 2006).

#### Graph theory analyses

Graphs consisted of recording sites (one per brain region) as nodes and links between regions. A link was drawn by thresholding based on significant recoding site–pair coherence in at least 80 % of the rats.

#### Multivariate Granger causality calculation

We used the MVGC toolbox(Barnett and Seth, 2014) to calculate Granger causality using the standard parameters for MVGC. Pairwise Granger causality magnitude was conditioned upon a one of the remaining recording sites and calculated once for each of the remaining sites. These conditioned magnitudes were then averaged to reduce spurious results due to global network activity. Statistical significance was assessed using a permutation test (× 1000, p < 0.01).

#### Phase–amplitude coupling analysis

We calculated the modulation index (MI) to quantify the degree to which the amplitude of gamma band oscillations are modulated by the phase of theta band oscillations. We used the method of Tort and colleagues(Tort et al., 2010b). The calculation was performed between recording sites that exhibited significant Granger causality. Following this method, we used surrogate data (see section on spectral analyses) to compute a surrogate distribution of MI values and subtract the threshold value at which the MI was significant for each phase–amplitude bin. As a result, any MI values reported in the paper are significant if they are greater than 0. For each of the 10 subjects, MI calculated for each phase–amplitude bin sparately for each pair of regions in each condition (baseline, stress, and coping). The across–subject average is presented as a co–modulogram.

#### Deep learning classifier

We used a built–in classifier from Matlab (ClassificationNeuronalNetwork) to predict experimental condition from coherence values calculated in 2–second bins during the first minute of the awake state for each condition. 834 and 672 observations were used to train the classifier and the remaining 20 % were used as test data). We computed the algorithm with a Bayesian optimization to obtain the best neuronal network possible based on our dataset (**Supplementary Table 1**). The fitcnet function was used to create the neural network classifier. The classifier was trained using a feedforward algorithm with a binary cross–entropy loss function. Accuracy was determined based on the number of times the prediction was correct across the total number of predictions. We evaluated the prediction accuracy using a surrogate distribution (× 1000) generated by shuffling the condition label (**Supplementary Fig. 2**). We assessed generalization to external data to avoid overtraining by holding–out data from one randomly selected rat from the training data set, and subsequently testing with held–out data (**Supplementary Fig. 3**).

### Statistical analyses

Only rats (n = 10) with correct electrode placement in all five regions of interest and with minimal signal artifacts during the entire experiment on at least two of the three electrodes were used for analyses. According to behavioral responses, following verification of the normality of the data, between-condition analyses were performed using a Wilcoxon test (struggling) or repeated– measures AVOVA and Neumann Keuls post hoc tests (all other variables). Permutation tests were Bonferroni corrected.

## Acknowledgements

The authors wish to thank Nathalie Gensbittel and Elouan Cosquer for technical help. This work was supported by the ANR (ANR–19–NEUR–0001–04 – NeuroMarKet) in the frame of the grant ERA– NET Neuron (LL), by the French government (PhD fellowship to LD), and by the Helsinki Institute of Life Sciences (HiLIFE) at the University of Helsinki (NT).

## Author contributions

Conceptualization: L.L., and L.D. Data curation: L.D. Formal Analysis: L.D., L.L., and N.K.T. Funding acquisition: L.L. Investigation: L.D., L.L., and K.H. Methodology: L.L., L.D., and N.K.T. Project administration: L.L. Supervision: L.L., and N.K.T. Visualization: L.D. Writing review and editing: L.D., N.K.T., and L.L.

