## Supplementary material for "Coping switches stress–evoked network disconnection to a new forebrain network state": supplentents

### Supplementary Information

#### Methods

##### Surgery

An analgesic (meloxicam; 1 mg/kg, s.c.) was administered prior to anesthesia induction (isoflurane; 4% in an induction box for 3 minutes; 1.5% for maintenance). A lidocaine/bupivacaine mix (2 mg/kg each, s.c.) was injected for local analgesia at the incision site. A bundle of three electrodes (0.004” PFA–coated tungsten, A–M Systems) were implanted in the right hemisphere in each region. Stereotaxic coordinates (mm) anteroposterior (AP) from Bregma, mediolateral (ML) from the midline of the sagittal sinus, and dorsoventral (DV) from the dura were as follows: AP = + 2.6, ML = 0.7, DV = – 3 (PRL); AP = + 0.5, ML = 0.7, DV = – 2 (ACC); AP = – 2.5, ML = 4.7, DV = – 6.8 (BLA); AP = – 3.5, ML = 0.7, DV = – 4.4 (LHb); and AP = – 3.8, ML = 2.7, DV = – 2.1 (CA1 region of the dHPC). A 2% DiI solution (Sigma–Aldrich) was applied at the tip of each electrode for post–mortem histological verification of their position. A reference screw was placed over left frontal cortex and a ground screw was places over the cerebellum. Once all electrodes were positioned, they were glued (C&B, Super–Bond) to the skull and skull screws that provided structural support. Wires were soldered to vias of an electrode interface board (EIB–16, Neuralynx). The wound margin was filled in with dental cement. Rats were placed in clean cages under a heating lamp until complete awakening. On the day following the surgery, rats received a second injection of meloxicam (1 mg/kg, s.c.). Following surgery, rats remained single housed. Experiments began following a 10–day recovery period.

##### Stress paradigm

First, over 3 consecutive days, for 3 hours/day, rats were habituated to being connected to the rotating commutator and to being moved to a different room and placed within a faraday cage); on the third day (designated as DAY 1 in **Fig. 1A**), neuronal activity was recorded as a baseline. On the following day, rats were exposed to repeated restraint stress consisting of two 10–minute restraint sessions performed 3 hours apart. Rats were connected to the recoding system for 1 hour prior to the first restraint. Recordings were performed during the 3 hours following each restraint. Restraint sessions were performed between 09:00 and 13:00 when CORT secretion, an indicator of the activity of the stress response axis, is minimal and stable[1]. The entire procedure was performed in the rat's home cage.

**Validation of the segregation of the signal into different vigilance states of interest**

We aimed to study stress and coping only in the awake state. Therefore, we defined vigilance states by a combination of the joint distribution of theta/delta power ratio in dHPC and delta/gamma power ratio in PRL, and the presence of body movement. Movement was measured by automated, markerless tracking of the rat's head from the video recordings using DeepLabCut (DLC) (2,3). Video was recorded (30 fps) using OBS Studio (Open Broadcaster Software 25.0.1). Training included 11 videos with 100 images and was created with the pre–trained network ResNet152. We used imaug (written by Alexander Jung; https://github.com/aleju/imgaug–doc) as an image augmenter and filtered the data using the median filter to remove outliers. We defined three vigilance states: active wake (AW), slow wave sleep (SWS), and rapid eye movement sleep (REM sleep). SWS was characterized by a high delta/gamma power ratio. AW and REM were characterized by a high theta/delta power ratio and low delta/gamma power ratio. REM was separated by lack of body movements. Other epochs that did not correspond to the above–mentioned characteristics were labeled quiet wake. Only AW data were analyzed.

**Assessment of the generalization of the feed–forward neuronal network**

We assessed whether the machine learning approach to predict experiment condition from coherency values could be generalized to held–out data. We trained the network on 9 out of 10 rats (**Fig. S2**, Test 1) and then repeated the procedure on the held out rat (**Fig. S3**, Test 2).

**Histology**

Rats were deeply anesthetized (pentobarbital, 120 mg/kg, i.p.). We performed intra–cardiac perfusion with 0.1 M PBS followed by 4 % paraformaldehyde (PFA; pH 7.4; 4°C). Brains were removed, post–fixed in 4 % PFA (4°C) for 48 hours, and then transferred into a 20 % sucrose solution (in 0.1 M PBS) at 4°C for 48 hours. Brains were frozen using isopentane (– 35°C) for 1 minute. Serial 40 µm–thick coronal sections were collected and mounted on gelatin–coated slides with a DAPI–fluoromount medium. DiI was visualized (Apotome.2, Zeiss®) to determine the location of the recording sites.

### Figures and analyses

#### Deep learning classifier settings

| *Parameters* | *AW* |
|---|---|
| *Layers size* | 68x197x7 |
| *Activations* | Tanh |
| *Output layer activation* | Softmax |
| *Solver* | LBFGS |

**Table S1**.

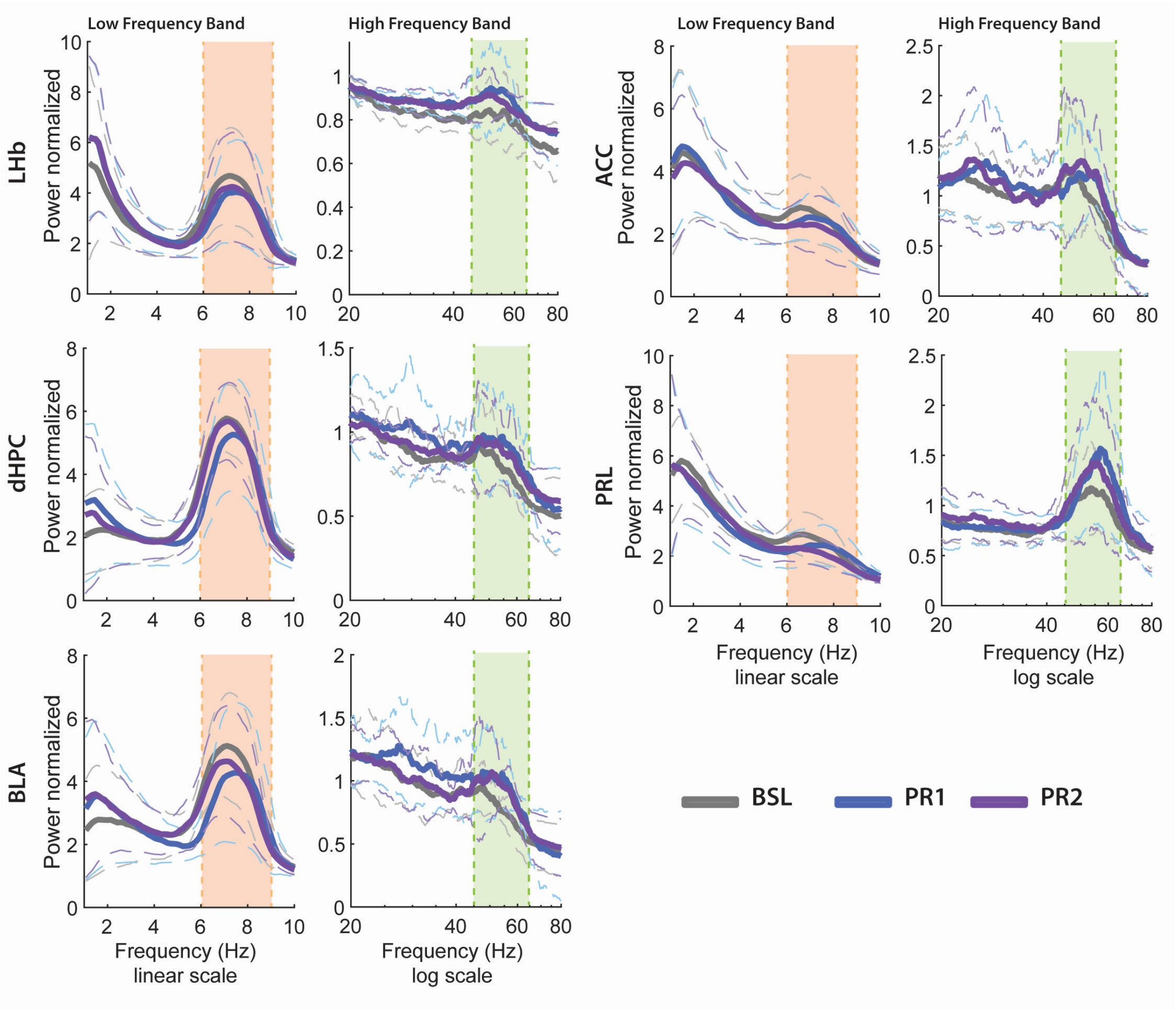


**Fig. S1**. Power spectrums of each brain region (LHb, ACC, dHPC, PRL, BLA) during the first AW min in the BSL (grey), PR1 (blue), and PR2 (purple) condition. Each power spectrum is separated into two parts according to low frequency band (1–10 Hz) with a linear scale for x axis, and high frequency band (20–80 Hz) with a logarithmic scale for x axis. Thick lines represent the standard deviation in the same color code than the given condition. The orange and green vertical columns represent delimitations of the frequency bands of interest, theta range (6–9 Hz) and gamma range (45–65 Hz) respectively.

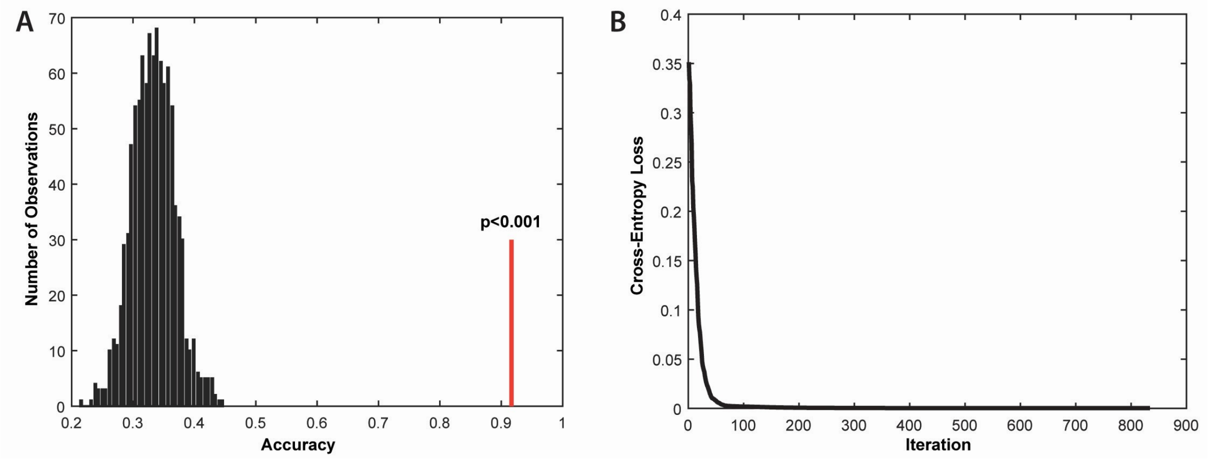


**Fig. S2**. Statistical validation and training overview of the classifier over the classes (BSL, PR1, PR2). The classes associated with each observation were shuffled (1000 repetitions) to assess that the results obtained were not spurious. We obtained a bootstrap distribution around 0.33 which corresponds to the fact that there were 3 classes. (**A**) Distribution of the bootstrap made on randomly relabeled observation (x 1000). The red line shows the location of the experimental observation. (**B**) Cross–Entropy Loss over the number of iterations during training of the classifier of the original observations representing how well the classifier performs.

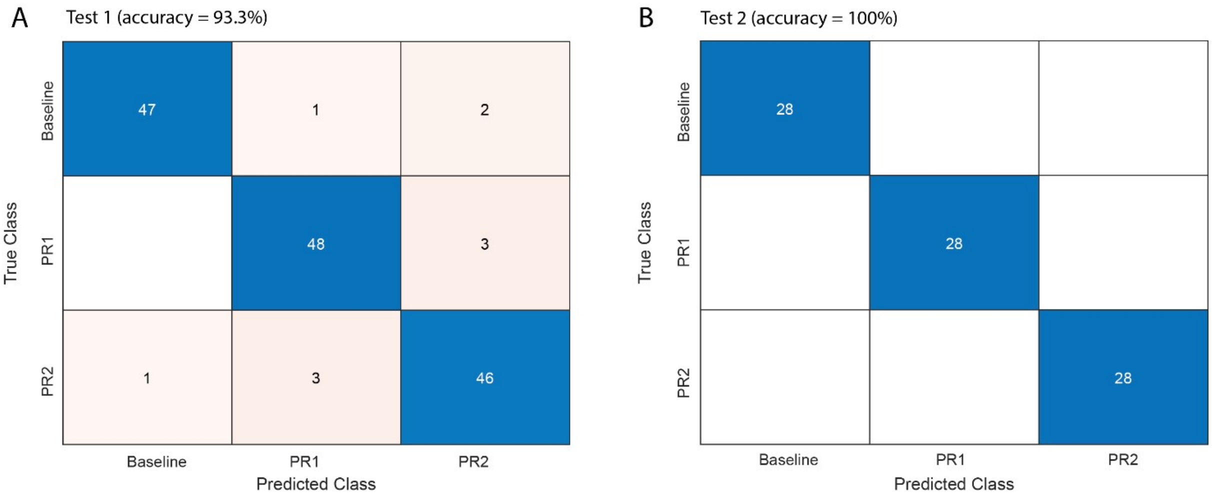


**Fig. S3**. Assessment of the generalization of the network to held out data. (**A**) Confusion chart of Test 1 of the neuronal network classifier based on 9 rats, which achieved accuracy of 93.3 %. (**B**) Confusion chart of Test 2, on the held–out rat. The accuracy was 100 % correct.
